# Inference of Cancer Drug Cross Resistance Using Only Single-Drug Exposure Data

**DOI:** 10.64898/2026.03.10.709822

**Authors:** Frederick J.H. Whiting, Ioanna Mavrommati, Ahmet Acar, Alvaro Ingles Russo Garces, Udai Banerji, Andrea Sottoriva, Rachael Natrajan, Trevor A. Graham

## Abstract

Combining drugs may prevent resistance in cancer treatment. However, the evolution of resistance to more than one drug simultaneously - cross resistance - remains a major obstacle to durable clinical response. Here, we show that we can identify the presence and strength of cross resistance using lineage tracing data from cell populations exposed only to each drug independently, removing the need for complex combinatorial treatment experiments. Simulations where the ground truth was known showed that the method accurately recovered the true underlying resistance dynamics. We applied the framework to breast cancer, lung cancer and lymphoma datasets, quantifying cross resistance across diverse therapy types. Our predictions align with biological expectations and reveal that even drugs with shared putative targets can exhibit variable cross resistance across pre-clinical models. Finally, we used inferred cross resistance strengths to evaluate treatment strategies, predicting that high cross resistance contributes to the limited efficacy of switching between CDK4/6 inhibitors. By enabling the identification of cross resistance without requiring multi-drug exposure experiments, our framework provides a method for prioritising drug combinations and screening for novel resistance-minimising drug targets.

## 1 Introduction

The evolution of drug resistance continues to limit the effective treatment of cancers. One strategy to tackle resistance is to combine different drugs to minimise the probability of resistance (Al-Lazikani, Banerji, and Workman 2012; Misale et al. 2015; Vasan, Baselga, and Hyman 2019; Plana, Adam C. Palmer, and Peter K. Sorger 2022). Molecular changes that render cells resistant to both treatments simultaneously are assumed to be more difficult to evolve and therefore occur less frequently than changes that provide resistance to either drug alone.

Cross resistance occurs when a resistance mechanism that confers resistance to the first drug simultaneously confers resistance to the second drug. Cross resistance can involve a diverse array of possible mechanisms that depend on the drugs in question. These include, among others: the over-expression of efflux pumps (Patch et al. 2015; Robey et al. 2018; Giddings et al. 2021; Wolf et al. 2025); the restoration of DNA-damage repair pathways (for example, homologous recombination) (Edwards et al. 2008; Pettitt et al. 2020); the alteration of the binding site of a shared target of multiple drugs (Yasuda, Kobayashi, and D. B. Costa 2012); and activation of bypass/downstream signalling for drugs targeting a shared pathway (Wee et al. 2009; C. Costa et al. 2020; Perurena, Situ, and Cichowski 2024). As with resistance to a single drug, the diverse array of possible mechanisms and the complex dynamics by which they emerge in cancer make identifying the phenotypes responsible challenging. There is now a wealth of evidence that resistant mechanisms can also be obtained via non-genetic means (Shaffer et al. 2017; White et al. 2025), compounding this challenge.

Measuring cross resistance is harder than measuring drug ‘interactions’ (differences in response due to antagonistic, additive or synergistic effects). These interactions measure the short-term response of a population and only require the measurement of viability following treatment with both drugs simultaneously. However, they contain no information on the durability of response to drug combinations. For example, a combination could be highly synergistic in the short-term but lead to rapid population regrowth in the medium-term due to strong cross resistance.

Statistical models such as ‘Bliss Independence’ (Bliss 1939) quantify the level of synergy by calculating the excess kill fraction above the expectation under drug additivity. Because resistant cells are typically rare prior to treatment, the measured response at concentrations that exert a strong selective pressure is dominated by the behaviour of the predominantly sensitive population. In contrast to these interactions, cross resistance is the product of a mechanism which is typically rare or absent. Its associated dynamics are therefore an emergent property of the population which requires long-term treatment exposure.

To measure cross resistance, a common approach is to generate cell populations that are resistant to each drug, usually via long-term exposure to treatment (Bhang et al. 2015; Dalin et al. 2022; Russo et al. 2022; Schaff et al. 2024). The resistant cells are subsequently exposed to concentrations of the opposing drug to measure the level of cross-resistance (Elms et al. 2025). One limitation of this approach is the lack of mechanistic insight it offers. For example, it would be unclear whether the cross-resistance that has evolved to each drug was due to a single, shared mechanism, two distinct mechanisms that each confer resistance to the opposite drug, or a combination of multiple competing mechanisms that exist in the population simultaneously. It would also be unclear whether resistance was the product of a stable, heritable change (genetic or epigenetic), or whether it occurred via transient non-genetic switching into a resistant cell phenotype. Clearly, these various possible evolutionary routes available to cancer cells pose difficulties for quantifying the dynamics of cross resistance and designing strategies to mitigate it (Gatenby et al. 2009; Dhawan et al. 2017; Maltas et al. 2024).

Here, we develop a framework to identify shared resistance mechanisms and to quantify the degree of cross resistance they confer using only data from single drug exposures as input. Our approach is agnostic to the molecular class of resistance mechanism (e.g. genetic or epigenetic) and instead detects cross resistance through analysis of clonal lineages, by looking for evidence that the same lineage is resistant to both drugs under single drug exposure. Importantly, defining cross resistance in terms of statistically associated resistant phenotypes within lineages enables the recovery of shared, heritable resistance mechanisms. We show the utility of our approach with examples from breast, lung and ovarian cancer and relate our findings to clinical trial results.

## 2 Results

### 2.1 A mathematical model of cross resistance

To quantify cross resistance evolution between cancer drugs, we developed a mathematical model that captures resistance to two drugs and is agnostic to the underlying molecular features causing resistance, thereby encompassing a diverse set of genetic and non-genetic evolutionary scenarios. Cross resistance is modelled as the statistical association between two cell phenotypes that confer resistance to one of two drugs. We build on our previous work that developed a framework that enables the quantification of resistance evolution to a single drug using cell population size changes and sequenced genetic barcodes from long-term resistance evolution experiments (Whiting et al. 2025). We aim to quantify cross resistance from lineage tracing experiments by leveraging the overlap of enriched cell lineages across drug treatments’ replicates. We reasoned that the enrichment of the same barcode subclones under different drugs is indicative of the presence of a shared, heritable resistance mechanism. Before constructing a model of cross resistance, we extend our previous two-phenotype framework, comprising sensitive (*S*) and resistant (*R*) cells, to accommodate resistance to two drugs (*Drug A* and *Drug B*). Cells can now occupy one of four phenotypic states: sensitive to both drugs (*S*), resistant to *Drug A* only (*R*_*A*_), resistant to *Drug B* only (*R*_*B*_), or resistant to both drugs simultaneously (*R*_*AB*_), referred to as double-resistant cells. We assume that cells transition unidirectionally from the sensitive phenotype to either single-resistant phenotype, *R*_*A*_ or *R*_*B*_, with probabilities *µ*_*A*_ and *µ*_*B*_ per cell division, respectively (Figure 1A). In the absence of cross resistance, the probability that a sensitive cell becomes double-resistant within a single division is given by the product of the single-resistance probabilities, *µ*_*AB*_ = *µ*_*A*_*µ*_*B*_. Each phenotype divides and dies with phenotype-specific rates *b*_*X*_ and *d*_*X*_, and drug treatment reduces cellular fitness through an increase in the effective death rate.

**Figure 1.**
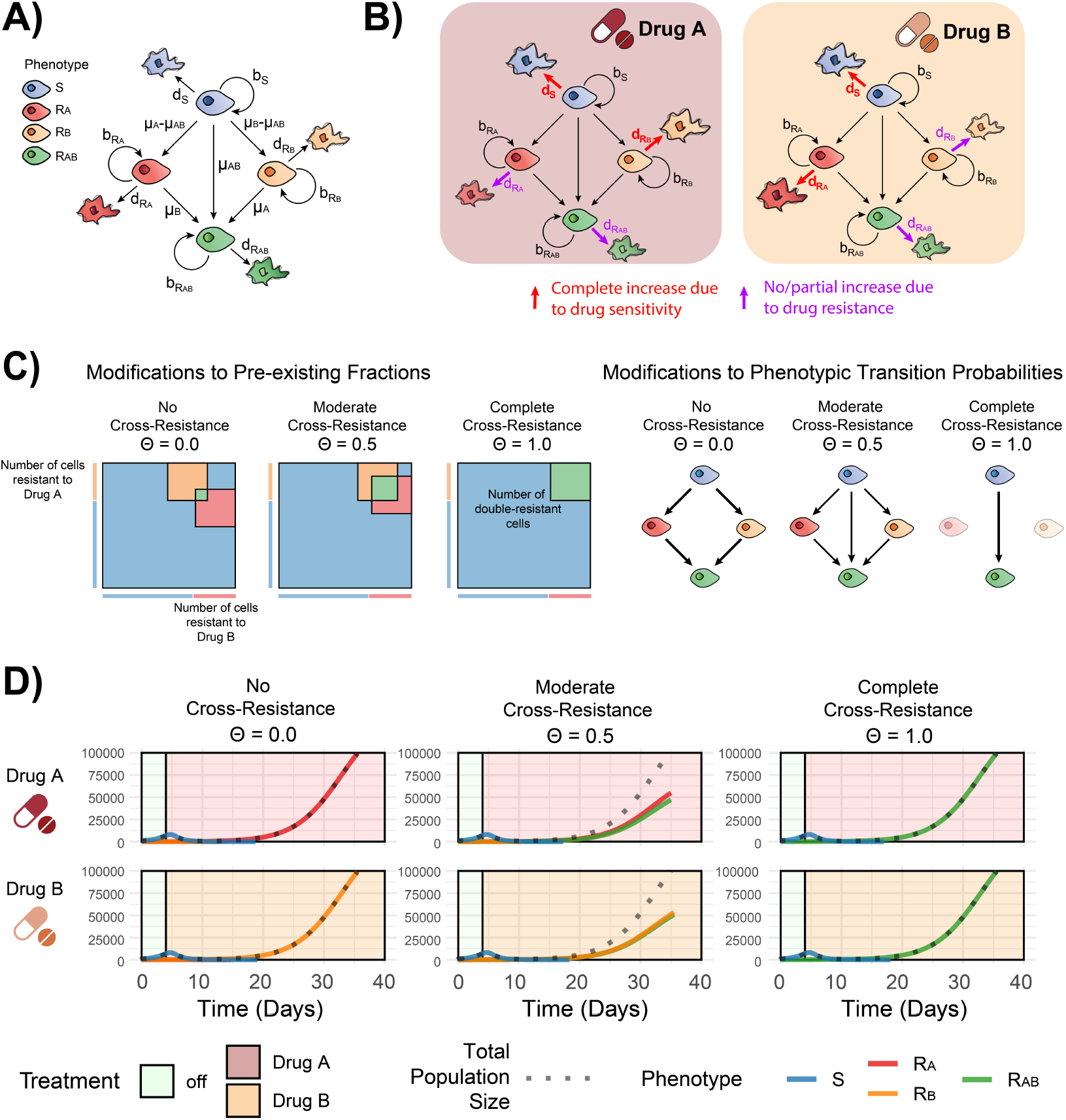
A two-drug resistance model can be modified to capture cross-resistance dynamics. **(A)** Phenotypes and phenotype-specific birth, death, and transition events in a model of resistance to two drugs. **(B)** Effects of each drug environment on phenotype-specific death rates. **(C)** Modifications to pre-existing phenotypic fractions (left) and phenotypic transition probabilities (right) under different strengths of cross resistance (*θ*). Border lines indicate the total fraction of cells resistant to each drug when considered independently, while shaded areas denote resistance occurring within the same cells (double-resistant) or in separate cells (single-resistant). **(D)** Illustrative phenotypic compartment dynamics under continuous treatment with each drug across varying strengths of cross resistance (*θ*).

In our model, each single-drug resistant phenotype confers no survival advantage under the alternative drug. Accordingly, cells resistant to only one drug have the same fitness as sensitive cells in the opposing drug environment (Figure 1B). Resistance phenotypes may be incomplete, where the drug-induced death rate in *Drug X* is reduced by a factor of *Ψ*_*X*_, relative to sensitive cells. This behaviour applies to both to singly resistant cells and to double-resistant cells when exposed to their corresponding drug.

Importantly, cells can be double-resistant without exhibiting a cross-resistant mechanism via the acquisition of resistance to each drug through two distinct independently acquired mechanisms. For example, if a cell contains two independent mutations, one that confers resistance to *Drug A* and the other to *Drug B*. We use the term ‘double-resistant’ to describe a phenotype, whereas ‘cross resistance’ describes a mechanism which dictates the association between these phenotypes. In the presence of a cross resistant mechanism, double-resistant cells occur at a higher frequency in a population than would be expected if resistance to each drug arose independently. For example, if now a single mutation conferred complete cross resistance to both drugs, resistant cells would exclusively exhibit the double-resistant phenotype. To model cross resistance, we therefore modify the pre-existing fraction of cells that exhibit each phenotype and the transition probabilities between each phenotype (Figure 1C). The strength of cross resistance (which we define as *θ*) increases the probability that the two resistant phenotypes are found together within cells. For example, under complete cross resistance (*θ* = 1.0) resistant cells exclusively pre-exist as, and transition from sensitive to, the double-resistant phenotype (assuming all other behaviours are the same for each resistant phenotype). Because some resistance mechanisms may be governed by molecular features that are incompletely heritable, such as the partial inheritance of epigenetic states during cell division, we allow *θ* to vary between 0.0 and 1.0.

We explored the model’s basic behaviour by examining how phenotypic compartments respond to treatment with either drug alone (Figure 1D). In the absence of cross resistance, each drug treatment selects for its respective resistant phenotype, killing cells that exhibit the remaining phenotypes. Increasing the strength of cross-resistance (*θ*) led to progressively larger fractions of double-resistant cells (*R*_*AB*_) under continuous monotherapy with either drug. Importantly, when considering only the total population size over time under each monotherapy, different cross resistance strengths are indistinguishable (Figure 1D).

### 2.2 Cell lineage information captures signatures of cross resistance in single drug exposure data

One way to infer the number of cross resistant cells in the population would be to re-treat the population with the alternate drug or subject them to combinations treatments. However, in these cases bulk population-level responses remain insufficient to resolve the underlying resistance mechanism dynamics. For example, survival under multiple drugs may arise from a single cross-resistant mechanism present within individual clones, or from the coexistence of distinct single-drug resistant clones within the population. We must therefore turn to alternative data types to resolve these dynamics.

Previously, we developed a framework that leveraged cell lineage (‘barcode’) information from long-term resistance evolution experiments to infer key evolutionary parameters (Whiting et al. 2025). Briefly, the approach relies on a shared mutual expansion period, following which closely related cells will share a barcode identity. The relationships between cell barcodes that survive treatment within and between experimental replicates subsequently enable the inference of behaviours such as the pre-existing fraction, relative fitness and transition rates between sensitive and resistant phenotypes. The development of this approach enabled us to quantify the resistance dynamics of cells exposed to a single drug.

We reasoned that this approach could be extended to experimental designs where populations of barcoded cells are exposed to different drugs in parallel. Specifically, we hypothesised that the relationships of cell lineages that survive within each drug condition should enable the quantification of drug-specific behaviours, whilst the relationships between drugs should capture the magnitude of cross resistance. As cells divide, shared cross resistance mechanisms that are heritable should reside within the same cell lineage. Consequently, exerting closely related cell populations to different drugs should select for the same cell lineages. Even in cases where the mechanism is only partially heritable, if a sufficiently short amount of time has passed before treatment begins, the majority of descendent cells will retain the shared mechanism. Because each barcode represents an individual subclone, this approach can be confident that the same mechanism is driving cross resistance to each drug, in contrast to cases where a cell population’s cross resistance is driven by polyclonal, competing mechanisms. Furthermore, this method precludes the difficulties around disentangling strong single drug resistance mechanisms from cross resistance mechanisms in situations where both drugs are given simultaneously.

We designed our model to be able to track individual cell lineages in a manner analogous to commonly adopted lineage tracing technologies. Specifically, in our agent-based model individual cells are assigned a unique (*in silico*) barcode when the simulation begins and share a mutual expansion period before being randomly sampled into replicate sub-populations (flasks) per drug condition (*Drug A* and *Drug B* - Figure 2A). Exploring the model’s behaviour with simulated data, we can also keep track of the size of each phenotypic compartment and the change in cell lineages during each timestep.

**Figure 2.**
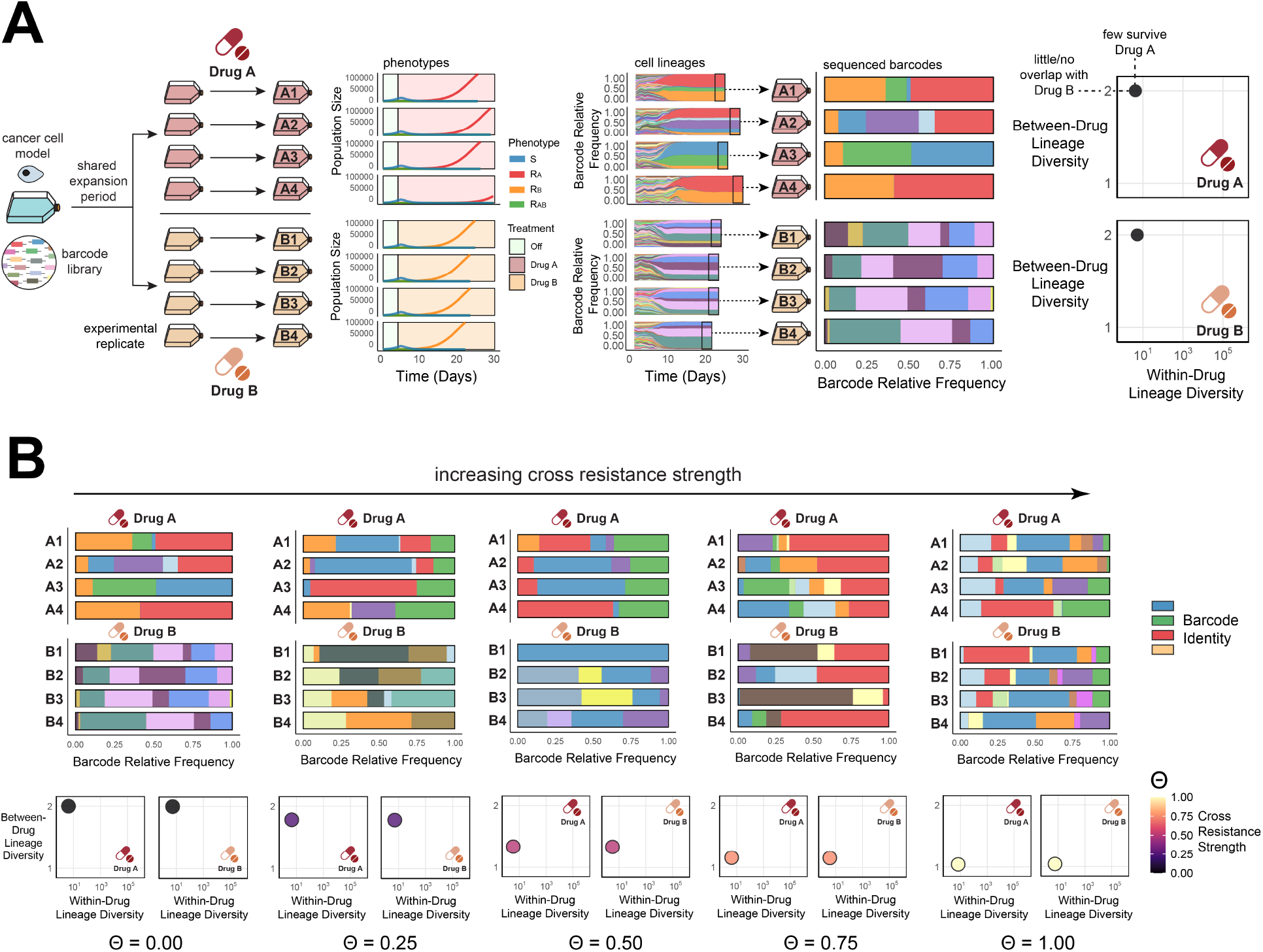
Cell lineage relationships generated from evolution experiments can capture the strength of cross resistance. **(A)** The experimental design and example simulated phenotypic compartment and cell barcode size change data from a lineage tracing experiment. Replicates are split into two conditions and treated with *Drug A* (A1-4) or *Drug B* (B1-4). Cell lineages are sampled at experimental endpoints and converted into within- and between-drug diversity statistics. **(B)** Between-drug diversity statistics track the cross resistance strength (*θ*) in simulated data where the true value of *θ* is known.

When we simulated resistance as a rare, stable phenotype, treatment gradually selects for the few, dominant lineages that exhibit the advantageous phenotype in each respective drug environment. In *in vitro* experiments, the full lineage histories are not observable and barcodes are only accessible when cells are harvested from the flask (Figure 2A). Despite this limitation, the resulting data can still consist of millions of unique barcode counts per replicate. By collapsing counts across replicates within each drug condition, we can estimate within- and between-drug lineage diversity using established diversity measures from ecology (Jost 2006; Chao and Jost 2015).

High between-drug diversity corresponds to low overlap in the surviving barcodes across drug conditions. Simulations using different strengths of cross resistance (encoded by different values of *θ*) showed that our model of cross resistance leaves a signature in this between-drug diversity statistic (Figure 2B). Importantly, because cross resistance does not impact the number of cells resistant to either drug alone, only the co-occurrence of the phenotypes, it has no impact on the within-drug diversity values (Extended Data Figure 1).

### 2.3 Cross resistance can be quantified using shared cell lineage data

We designed a statistical inference framework to quantify cross resistance using input data on cell population sizes and genetic barcode distributions.

We first quantify the resistance dynamics to each drug independently. Under our model formulation, cells resistant to *Drug B* (*R*_*B*_) do not alter their sensitivity to *Drug A* and therefore are indistinguishable from sensitive cells (*S*). We therefore treat *R*_*B*_ cells as sensitive when inferring within-drug parameters for *Drug A* and use our simpler model that tracks only sensitive (*S*) and resistant (*R*) compartments. An analogous simplification applies when analysing resistance to *Drug B*, where *R*_*A*_ cells are treated as sensitive. For each *Drug X* we use a likelihood-free simulation-based Bayesian inference framework (Methods) to independently estimate the within drug behaviours: the pre-existing fraction of resistant cells (*ρ*_*X*_), the probability of transition from sensitive to resistant per cell division (*µ*_*X*_), the effective drug concentration (*Dc*_*X*_), the strength of the resistant phenotype (*Ψ*_*X*_), and the drug accumulation and decay rate (*κ*_*X*_). Inference proceeds in two stages. First, we use a hybrid phenotypic compartment model that tracks only the sizes of the sensitive and resistant populations (*n*_*S*_ and *n*_*R*_) to exclude regions of parameter space inconsistent with the observed population size dynamics. We then fit to the cell barcode data using a more computationally intensive agent-based model that explicitly tracks individual cell lineages to estimate the full posterior distributions of the within-drug parameters (Figure 3A).

**Figure 3.**
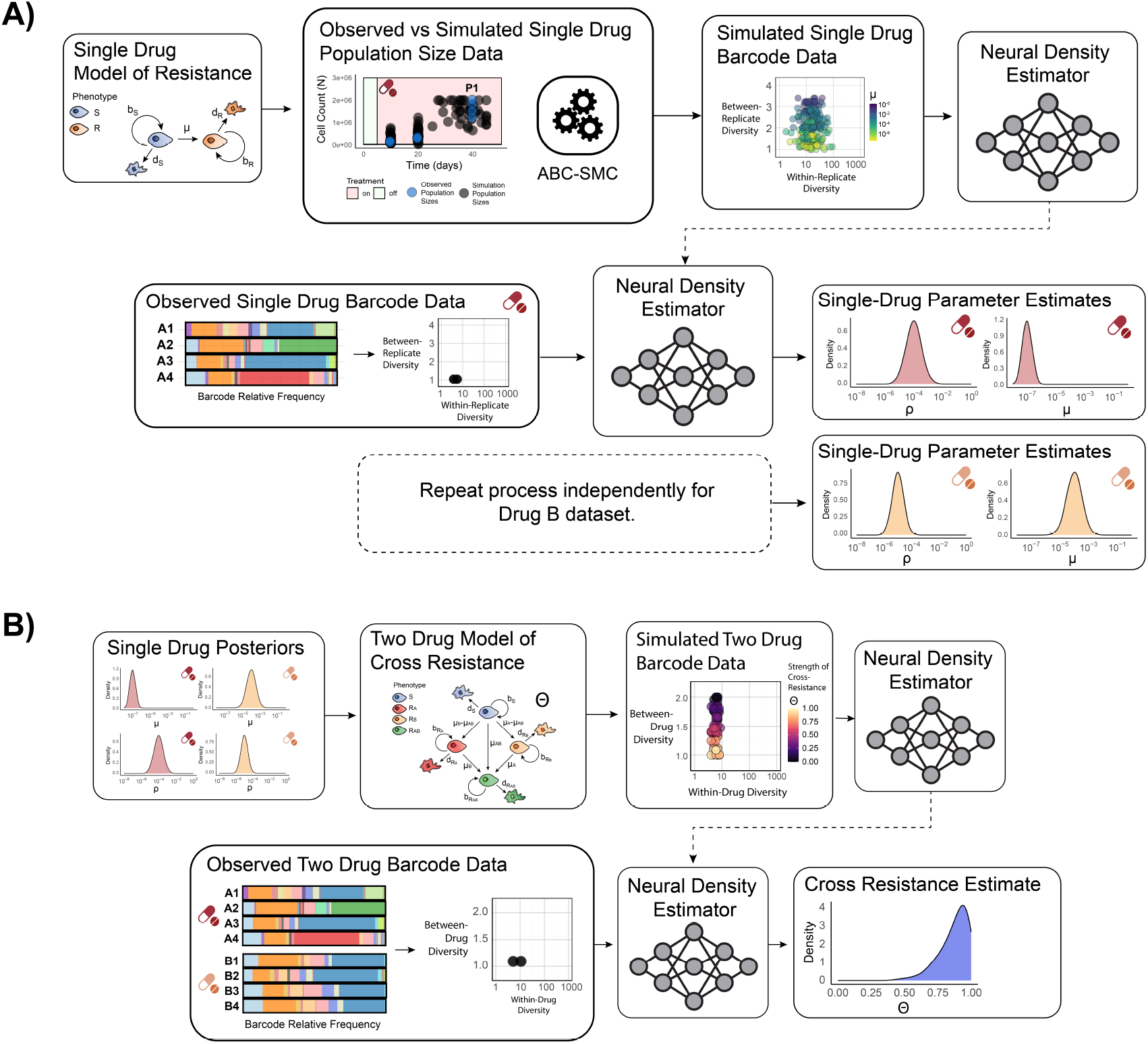
A joint framework to infer the evolutionary behaviours driving resistance to two drugs and the strength of cross resistance between them. **(A)** Workflow for estimating single-drug evolutionary parameters. Inference proceeds in two stages per drug. First, a hybrid phenotypic compartment model is fit to population size trajectories using ABC–SMC. Second, parameter sets retained by the ABC–SMC procedure are used to simulate the agent-based barcode model, where barcode distributions’ diversity statistics are provided to a neural posterior estimator (NPE). Observed data are passed through the trained NPE to obtain posterior distributions. A single illustrative parameter (*µ*) is shown in the ‘Simulated Single Drug Barcode Data’ panel. **(B)** Workflow for estimating the strength of cross resistance (*θ*). Parameter sets for each drug are drawn from their respective single-drug posteriors and combined with values of *θ* sampled from the prior. The two-drug model is simulated and barcode distributions are converted into between-drug diversity statistics. These statistics are provided to a NPE framework to obtain the posterior distribution of *θ*.

We infer the strength of cross resistance using the two-drug cross resistance model. To do this, we draw parameter sets for each drug independently from their respective within-drug posterior distributions and simulate the two-drug model across a range of cross-resistance strengths (*θ*). Inference is performed using a simulation-based inference (SBI) framework (Boelts et al. 2025), where *θ* is informed by the between-drug diversity statistics (Methods).

### 2.4 Validation of inference framework

We validated our inference framework using simulated data for which the true parameter values were known. We first considered a simple scenario of stable, pre-existing resistance, where the drug exerted a strong effect on sensitive cells and the resistant phenotype was complete (Figure 4). We assumed identical within-drug behaviour for both drugs, such that the pre-existing resistant fractions were equal (*ρ*_*A*_ = *ρ*_*B*_). Under these conditions, the framework accurately recovered the within-drug parameters and used the between-drug barcode relationships to distinguish between low, moderate, and high levels of cross resistance (Figure 4A–C). We repeated this procedure for a range of biologically plausible scenarios and found that we could recover the strength of cross resistance (*θ*) across a diverse set of evolutionary contexts (Extended Data Figure 2). Importantly, our framework only infers strong cross resistance when it is truly driving resistance to both drugs.

**Figure 4.**
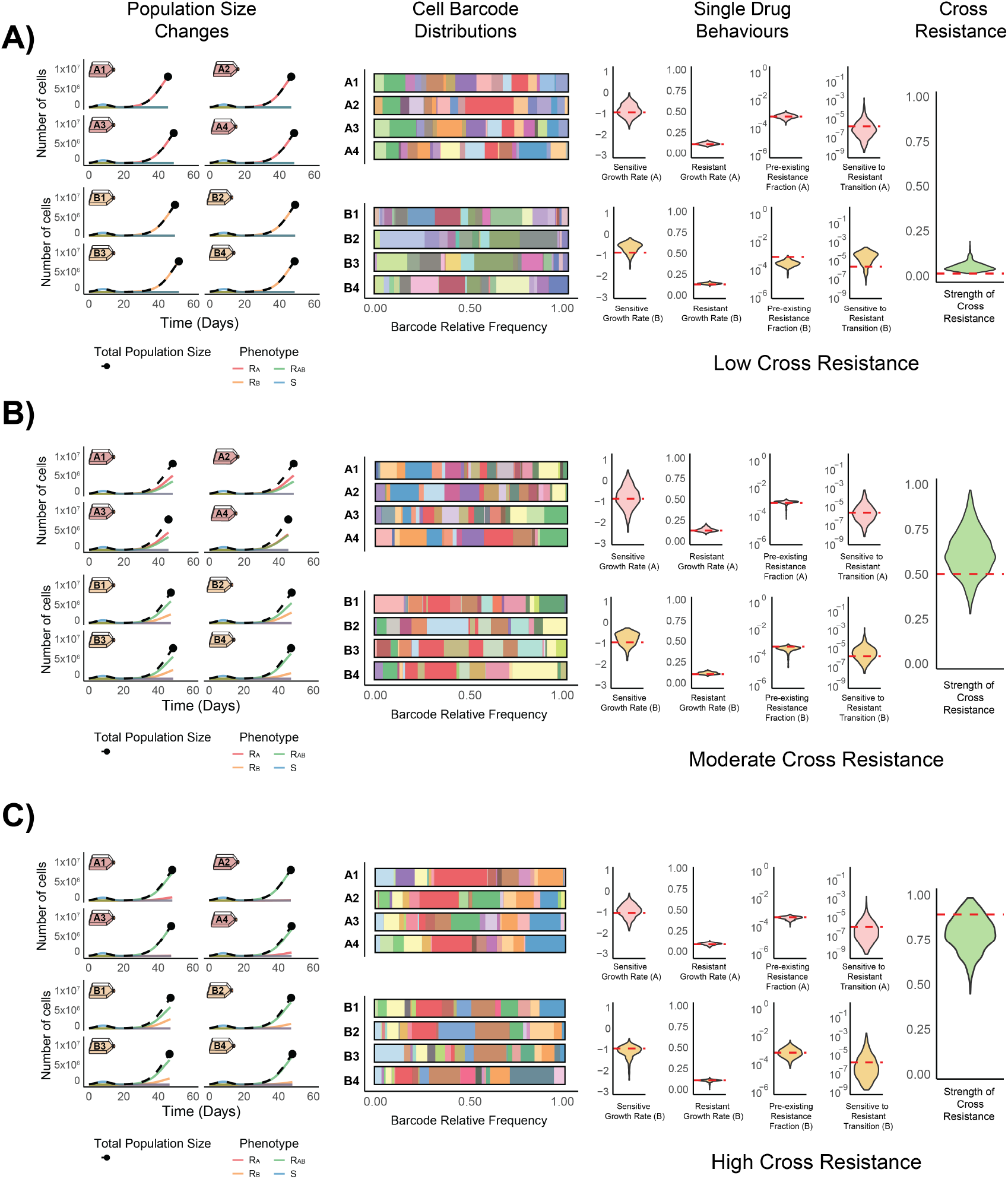
Cell lineage relationships enable the robust inference of known cross resistance strengths from synthetic data. Three synthetic datasets generated with low **(A)**, moderate **(B)** and high **(C)** strengths of cross resistance (*θ*). Total cell population sizes (points) and the cell barcode distributions used for single drug behaviour and cross resistance estimation. Inferred posterior distributions (shaded densities) and ground-truth parameter values (red dashed lines) are shown.

We used our framework to explore the conditions under which cross resistance could be reliably inferred. We found that scenarios in which resistance to each drug was more frequent led to more robust estimates of the cross resistance strength, *θ* (Extended Data Figure 2): when resistant cells were rare, only a small number of cell lineages contributed to resistance in each condition. This increased stochastic variation in which resistant lineages were sampled into each replicate flask, leading to greater variability in the observed overlap of resistant lineages between drug conditions. Because our approach relies on detecting shared resistant lineages between drug conditions, this sampling noise resulted in less confident estimates of *θ*.

The degree of asymmetry between the resistant phenotypes to each drug was another factor that influences our ability to infer *θ*. For example, holding all other parameters constant between *Drug A* and *Drug B*, estimates of *θ* became less reliable as the pre-existing fraction of resistance to one drug increasingly outnumbered the other (Extended Data Figure 3A). That is, inference performance deteriorated as asymmetry between *ρ*_*A*_ and *ρ*_*B*_ increased. This effect has a simple statistical explanation. In our model, the pre-existing fraction of double-resistant cells is given by *ρ*_*AB*_ = (1 *− θ*)(*ρ*_*A*_*ρ*_*B*_) + *θ* min(*ρ*_*A*_, *ρ*_*B*_). Under complete cross resistance (*θ* = 1.0) this reduces to *ρ*_*AB*_ = min(*ρ*_*A*_, *ρ*_*B*_). As one resistant phenotype becomes more common than the other, the double-resistant cells constitute an increasingly smaller fraction of the total resistant population. Because shared cell lineages correspond to cells that exhibit the double-resistant phenotype, greater asymmetry reduces the number of lineages that are selected for under both treatments. This weakens the signal captured by the between-drug diversity statistics and leads to poorer recovery of *θ*.

This reasoning extends beyond pre-existing resistance. We found that the same limitation arose for other model features that influence the number of cells surviving each treatment, for example asymmetry in the sensitive-to-resistant transition probabilities (*µ*_*X*_) and the effective strength the drug exerts on the cell phenotypes (*Dc*_*X*_) (Extended Data Figure 3B and 3C). In all cases, increasing asymmetry reduced the fraction of lineages jointly enriched across drug conditions, limiting the information available to infer cross resistance.

Taken together, these results show that our framework can robustly recover cross resistance across a broad range of evolutionary scenarios. Where sufficient numbers of shared double-resistant cell lineages exist, we consistently infer the presence and strength of cross resistance, while correctly identifying cases where cross resistance is weak or absent.

### 2.5 The quantification of cross resistance across diverse disease areas and therapies

We applied our framework to experimental data from long-term resistance evolution experiments spanning three disease contexts and eight cancer drugs. For all experimental studies considered here, we used a single passage of cell population size observations together with the sequenced barcode distributions from each treatment condition’s replicates. Parameters fixed by the experimental design including treatment schedules and sampling structure were specified beforehand (Methods). All evolutionary parameters were inferred from the data using our inference procedure (Figure 3).

#### 2.5.1 Breast cancer

We analysed previously reported data that evolved resistance to two CDK4/6 inhibitors (abemaciclib and palbociclib) in three ER+ breast cancer cell models (BT474, MCF7, and T47D) (Mavrommati et al. 2025). Across all three cell models, there was evidence of cross resistance between the two CDK4/6 inhibitors, although the inferred strength varied by model (Figure 5A). In BT474 cells, we inferred strong cross resistance, with a median estimate of *θ* = 0.866 (95% credible interval: 0.650–0.988) and in T47D cells we inferred moderately strong cross resistance with a median value of *θ* = 0.689 (95% credible interval: 0.399–0.945). Cross resistance in MCF7 cells was inferred to be lower with a median of *θ* = 0.365 (95% credible interval: 0.056–0.711) (Figure 5A). Inferred within-drug parameters revealed cell-model-dependent asymmetries between the two treatments (Extended Data Figure 4A), but these were not sufficiently pronounced to prevent the inference of strong cross resistance.

**Figure 5.**
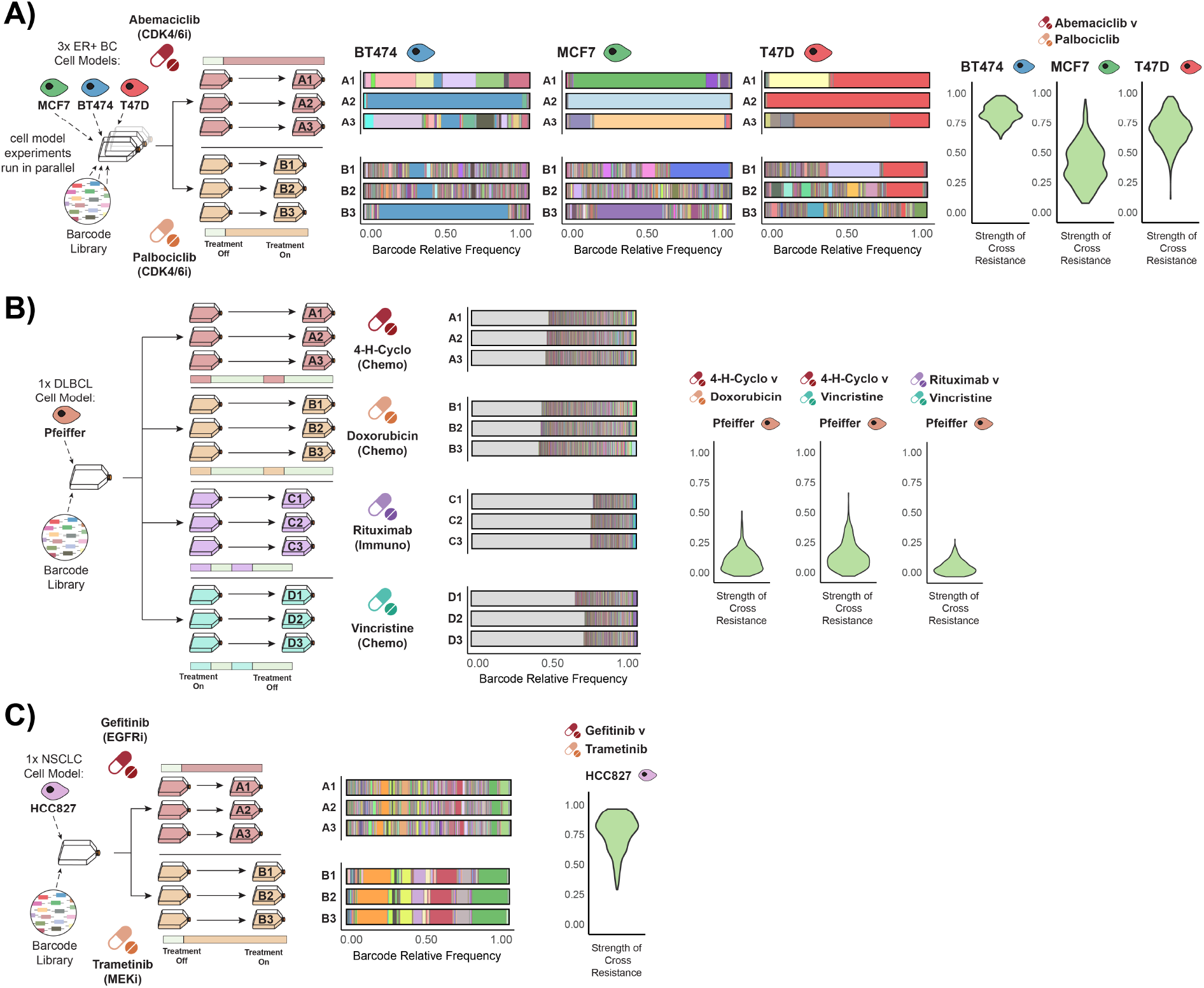
Cross resistance varies by disease and cell model across a diverse set of treatments. Experimental design schematics, sequenced barcode distributions and cross resistance strength estimates for three datasets: **(A)** two CDK4/6 inhibitors (Abemaciclib and Palbociclib) in three ER+ breast cancer cell models (BT474, MCF7 and T47D), **(B)** 4-hydroperoxy-cyclophosphamide (4-H-Cyclo), Doxorubicin, Rituximab, and Vincristine in a diffuse large B-cell lymphoma (DLBCL) cell model, **(C)** an EGFR inhibitor (Gefitinib) and a MEK inhibitor (Trametinib) in a non-small cell lung cancer (NSCLC) cell model. Colours denoting unique cell barcode lineages are only consistent across replicates within each cell model. Grey areas indicate barcodes with a relative frequency *<* 10^*−*3^.

Notably, in the study from which these data were derived (Mavrommati et al. 2025), T47D cells were shown to exhibit RB1 loss following resistance evolution. Altogether, in T47D cells our framework predicts a stable, pre-existing mechanism that exhibits strong cross resistance. Loss of RB function is a well-established mechanism of resistance to CDK4/6 inhibition (Herrera-Abreu et al. 2016; Condorelli et al. 2018) and has previously been reported as driving resistance in this cell model (Algethami et al. 2025; Migliaccio et al. 2025).

#### 2.5.2. Lymphoma

We applied the framework to a dataset in which a diffuse large B-cell lymphoma (DLBCL) cell model (Pfeiffer cells) was exposed to the constituent drugs of the R-CHOP regimen (Adam C Palmer, Chidley, and Peter K Sorger 2019). We focused on four drugs: 4-hydroperoxy-cyclophosphamide (4-H-Cyclo), Doxorubicin, Rituximab, and Vincristine. In this experiment, the cell population had been subjected to random mutagenesis prior to treatment. Combined with the short, intermittent treatment schedule, this resulted in substantial barcode diversity being retained across all conditions (Figure 5B). Our within-drug inference captured these dynamics, estimating relatively large pre-existing fractions of resistance, particularly in the Rituximab and Vincristine conditions where lineage diversity was highest (Extended Data Figure 4B).

Because the experimental design involved four drugs, we inferred cross resistance using pairwise comparisons between treatment conditions. We selected three comparisons to illustrate the approach: 4-H-Cyclo versus Doxorubicin (4-D), 4-H-Cyclo versus Vincristine (4-V), and Rituximab versus Vincristine (R-V). Across all three comparisons, we inferred low levels of cross resistance: 4-D (*θ* = 0.109, 95% credible interval: 0.011–0.322), 4-V (*θ* = 0.137, 95% credible interval: 0.012–0.399), and R-V (*θ* = 0.064, 95% credible interval: 0.005–0.201) (Figure 5B).

Our results indicating low cross resistance across the RCHOP components in DLBCL are consistent with the original study (Adam C Palmer, Chidley, and Peter K Sorger 2019) and supports their interpretation that limited overlap in resistance mechanisms contributes to the curative efficacy of this combination.

#### 2.5.3. Lung cancer

Finally, we applied the framework to a dataset generated by exposing a non-small cell lung cancer (NSCLC) cell model to two MAPK-pathway-targeting therapies: gefitinib (an EGFR inhibitor) and trametinib (a MEK inhibitor) (Acar et al. 2020). Similar to our results supporting strong cross resistance to two CDK4/6 inhibitors, we inferred high levels of cross resistance between these two MAPK-targeting agents (*θ* = 0.806, 95% credible interval: 0.381–0.987, Figure 5C) despite modest asymmetries in the inferred within-drug parameters (Extended Data Figure 4C).

In the original study (Acar et al. 2020), analysis revealed a substantial fraction of barcodes that were enriched following independent exposure to EGFR and MEK inhibition, which were described as ‘double-resistant clones’. Our framework naturally interprets this overlap as evidence of cross resistance between the two drugs. Cross resistance is biologically plausible given that EGFR and MEK inhibitors act vertically within the MAPK pathway and resistance to either can converge on the same mechanism (Tricker et al. 2015; Oddo et al. 2016).

### 2.6 Cross resistance information can inform treatment switching strategies

We explored how estimates of cross resistance (*θ*) could inform treatment switching strategies. The efficacy of a second-line therapy will depend on the degree of cross resistance between it and the first-line drug, since strong cross resistance implies that selection for resistance to the first drug may have already enriched resistance to the second. Using our inferred within- and between-drug parameters, we designed a simulation-based approach to quantify the potential benefit of treatment switching (Extended Data Figure 5A). In this framework, a synthetic patient is split into two simulation branches when the tumour reaches an initial progression threshold (*N*_*p*_) under first-line treatment (at time *T*_*A*_ for *Drug A*, for example): in one branch treatment continues unchanged, while in the other treatment switches to a second-line drug. Treatment then proceeds until a clinical failure threshold is reached (*N*_*cf*_), and the difference in time between the branches (Δ*T*_*AB*_ = *T*_*AA*_ *− T*_*AB*_, for example) defines the clinical benefit of switching (Methods).

Following confirmation that our inference framework could robustly recover the true dynamics of switching regimens (Extended Data Figure 6), we examined how variation in within-drug behaviours influences switching efficacy. We fixed the parameters governing response to *Drug A*, and then jointly sampled parameters controlling response to *Drug B* together with a range of cross-resistance strengths (*θ*). For each parameter set, we quantified the absolute time gained or lost following treatment switching (Methods).

We found that the efficacy of treatment switching depended strongly on asymmetry in within-drug behaviours, in addition to the strength of cross resistance (Extended Data Figure 5C). Across the parameter ranges explored, asymmetries in resistant phenotype dynamics often had a greater overall impact on switching benefit than *θ* itself. Nonetheless, increasing cross resistance consistently reduced the clinical benefit of switching (lower Δ*T*_*AB*_ and Δ*T*_*BA*_) across a wide range of conditions. For example, when the pre-existing resistant fractions were approximately symmetric, *θ* controlled the magnitude of benefit gained following treatment switching (Extended Data Figure 5C, *ρ*_*B*_ = 10^*−*3^).

Taken together, these results demonstrate that a quantitative assessment of treatment switching strategies requires knowledge of both within-drug evolutionary dynamics and the between-drug cross resistance strength. Whilst the strength of cross resistance constrains the potential benefit of switching, it is not sufficient alone to predict optimal strategies without accounting for possible asymmetries in drug-specific resistance behaviours.

Finally, we extended this analysis to synthetic clinical trials using our inferred parameter distributions from the three ER+ breast cancer cell models dataset. Each posterior distribution defined for a given cell model and drug comparison can be interpreted as a ‘patient archetype’ capturing underlying biological features of drug resistance. By sampling from these distributions, simulating treatment and measuring time to progression (TTP), we predicted archetype-specific responses to treatment switching in a clinical trial setting (Figure 6A; Methods).

**Figure 6.**
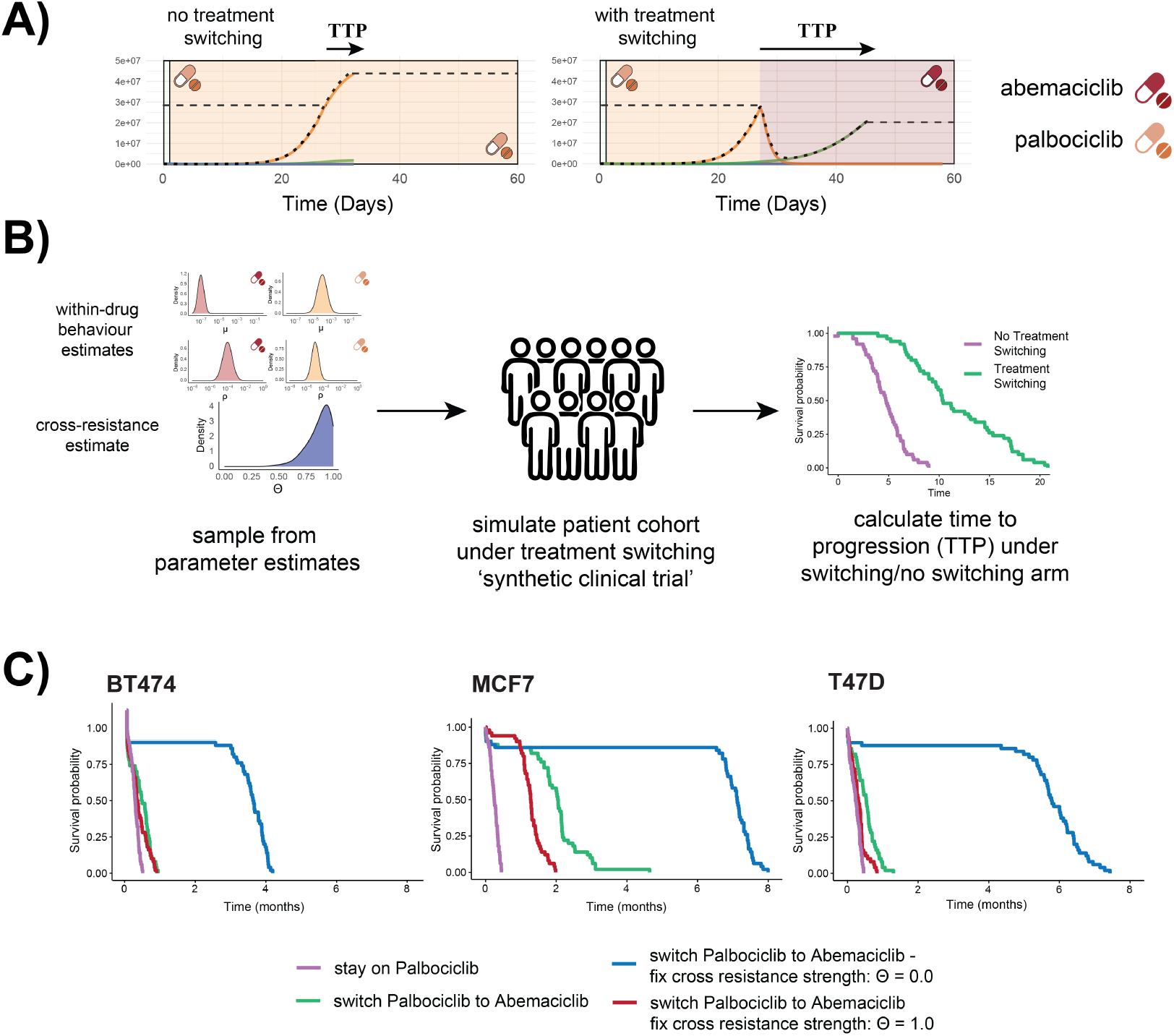
Testing clinical treatment switching under different cross-resistance scenarios. **(A)** Time to progression (TTP) is calculated under a no-switch control arm and a treatment-switching arm following progression on first-line therapy (left dashed line; *N*_*p*_). After progression, tumour burden is monitored every 7 days and subsequent progression is defined as the first time the population exceeds 0.5 *× N*_*p*_. **(B)** Workflow for generating synthetic patient cohorts using posterior estimates of within-drug parameters and cross resistance. For each posterior draw, TTP is calculated under both continued treatment and treatment switching. **(C)** Estimated resistance parameters for Abemaciclib and Palbociclib in ER+ breast cancer cell models are used to compare TTP distributions across three patient archetypes. Treatment switching (green) vs no switching (purple) is evaluated alongside theoretical limits of no cross resistance (*θ* = 0.0, blue) and complete cross resistance (*θ* = 1.0, red).

A key advantage of our mechanistic framework is that individual parameters can be fixed, allowing their specific contributions to clinical outcomes to be isolated. In particular, by fixing the cross resistance strength (*θ*) while continuing to sample all within-drug parameters, we defined a cross resistance window corresponding to the minimum and maximum predicted TTP distributions under *θ* = 0 and *θ* = 1, respectively. This allowed us to directly assess the extent to which the predicted benefit or failure of treatment switching could be attributed to cross resistance itself, rather than to asymmetries in drug-specific resistance dynamics.

Using our three ER+ breast cancer model archetypes, we simulated a synthetic clinical trial comparing a palbociclib-to-abemaciclib switching arm with a continuous palbociclib control arm (Figure 6B). This switching direction was chosen given recent clinical trials exploring similar conditions, although typically in combination with endocrine therapies (Kalinsky et al. 2025). Across all three archetypes, we observed little to no improvement in TTP for the BT474 and T47D models, and only a moderate benefit for MCF7. In all cases, the predicted TTP distributions lay close to the lower bound of the cross-resistance window, indicating that cross resistance was a major contributor to the limited clinical benefit of CDK4/6i switching.

## 3 Discussion

We developed a mechanistically tractable framework for quantifying cross resistance that models cross resistance as the statistical association between resistant mechanisms within evolving cell lineages. We find that the independent selection for the same resistance mechanism leaves a signature in the form of enrichment of cell lineages across independent single drug exposures. Once the behaviour to each single drug has been quantified, we show that the shared clonal lineage information can be used to infer the strength of cross resistance across a range of evolutionary scenarios.

Identifying strong cross resistance is particularly valuable because it points to convergent evolutionary constraints. When resistance to multiple drugs repeatedly arises through the same mechanism, this implies dependencies that may be therapeutically targetable. More generally, such findings can inform the prioritisation of drug combination strategies that explicitly avoid shared evolutionary escape routes. We predict our approach will be most valuable when identifying non-intuitive cross-resistance relationships, where its systematic approach across diverse drug classes could reveal unexpected evolutionary relationships that would be difficult to anticipate.

An advantage of our approach is that it can identify the presence of cross resistance mechanisms only using data routinely generated in long-term experimental evolution studies, namely total cell population size changes and barcode lineage distributions. Moreover, because inference is based on comparisons between independent drug exposure experiments initiated from the same ancestral barcode pool, it is not necessary to perform multiple, arduous cross-exposure experiments. Any pair of drugs for which evolution experiments have been performed from a common barcoded cell population can be compared. This enables the analysis of cross resistance at scale.

Recent efforts have focussed on identifying synergistic drug combinations that can make use of lower concentrations of each drug whilst maintaining a high selective pressure (Jaaks et al. 2022). One obvious advantage of synergy is the potential reduction in off-target patient toxicity effects. However, evidence suggests that in some cases synergistic combinations might lead to higher rates of cross resistance (Mason-Osann et al. 2024). This occurs because synergism is by definition reliant on the interaction between drugs. For example, the dual inhibition of the MAPK and PI3K pathways, where cross-talk between the cascades means blockade of one enhances the effect of blocking the other (Murali et al. 2021). Alongside benchmarking drugs by their propensity for intra-drug resistance and strength of inter-drug cross resistance, our approach provides a framework with which the relationship between synergy and cross resistance can be tested quantitatively, with implications for finding drug combinations that maximise clinical benefit.

Our model necessarily makes a number of simplifying assumptions. We neglect spatial structure, assume well-mixed populations and focus on a single dominant resistance mechanism per drug. This choice is motivated by our implementation in an *in vitro* setting and by the expectation that under strong, continuous selection the fittest resistance mechanism will tend to dominate the population dynamics. In such cases, we argue the behaviour of the population is well approximated by tracking the phenotypes that determine the ‘winning clone’. However, in scenarios where multiple resistance mechanisms coexist over long timescales and confer qualitatively different growth or switching behaviours, a single-mechanism approximation may fail to fully capture the dynamics. We note that this is more likely for patients’ solid tumours, where the spatial separation of expanding clones could lead to multiple mechanisms emerging during treatment (Awad et al. 2021; Harvey-Jones et al. 2024). Developing the framework to explicitly accommodate multiple competing mechanisms would be a natural extension.

We present an extensible framework to infer drug cross resistance at scale without requiring sequential resistance evolution or cross exposure experiments. This enables the simultaneous benchmarking of drugs, identification of shared resistance mechanisms and design of treatment sequences that delay resistance emergence.

## Acknowledgements

This study was principally funded by Cancer Research UK by the programme grant DRCNPG-May21 100001 to T.G. T.G. and U.B. acknowledge funding from the CRUK convergence science centre and Biomedical Research Centre. R.N. receives funding from the Breast Cancer Now’s Catalyst Programme (Grant 2017JulyPCC005) which is supported by funding from Pfizer; and by Programme Grants from Breast Cancer Now as part of Programme Funding to the Breast Cancer Now Toby Robins Research Centre. A.S. is supported by the Associazione Italiana per la Ricerca contro il Cancro (AIRC) (28961) and by the ERC Consolidator Award (101125077). A.A. is supported by the National Outstanding Researchers Program administered by The Scientific and Technological Research Council of Türkiye (TÜBİTAK) (123C588).

## 6 Methods

### 6.1 Phenotypic model of single-drug resistance

We previously described a phenotypic compartment model in which cells exist in one of two mutually exclusive states: sensitive (*S*) or resistant (*R*) (Whiting et al. 2025). The numbers of sensitive and resistant cells at time *t* are *n*_*S*_(*t*) and *n*_*R*_(*t*), respectively, with total population size *N* (*t*) = *n*_*S*_(*t*) + *n*_*R*_(*t*). The initial resistant fraction is *ρ* = *n*_*R*_(0)*/N* (0).

Population growth is logistic with carrying capacity *C*. Each phenotype has birth and death rates that depend on the current effective drug concentration. Phenotypic transitions are coupled to cell division: sensitive cells acquire resistance with probability *µ*per division and in this model resistance is assumed to be irreversible.

Drug treatment is incorporated through a time-dependent effective concentration *D*(*t*) = *Dc, γ*(*t*), where *Dc* controls drug strength and *γ*(*t*) *∈* [0, 1] follows an uptake–decay process during treatment on/off windows. Drug exposure increases cell death rates. Sensitive cells experience

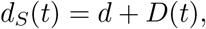

whereas resistant cells experience reduced drug-induced mortality,

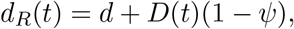

with *Ψ ∈* [0, 1] controlling the ‘strength’ of the resistant phenotype.

### 6.2 Two-drug extension

To model resistance to two drugs, we extended the framework to four mutually exclusive phenotypic states: sensitive (*S*), resistant to Drug A only (*R*_*A*_), resistant to Drug B only (*R*_*B*_), and resistant to both drugs (*R*_*AB*_). The total population size at time *t* is

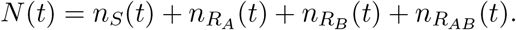

Cells transition to the resistant phenotype to Drug A or Drug B during cell division with probabilities *µ*_*A*_ and *µ*_*B*_ per cell division, respectively. In the absence of cross resistance, acquisition is independent and double resistance arises with probability *µ*_*A*_*µ*_*B*_. Resistance is assumed to be irreversible.

Drug-dependent death rates depend jointly on the phenotype and drug environment.

Under Drug A treatment,

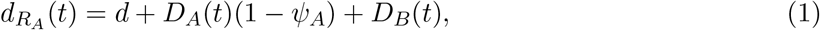

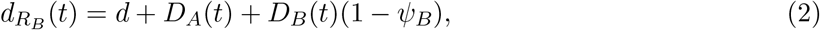

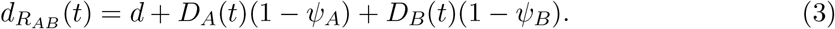

The model allows exposure to one or both drugs, although in this study we only explore scenarios where drugs were applied individually.

### 6.3 Modelling cross resistance

Cross resistance is defined as the statistical association between resistance phenotypes exhibited by individual cells. Under Law independence (Law 1956), the expected fraction of double-resistant cells equals the product of single-drug resistance probabilities,

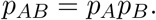

To model varying levels of cross resistance, we introduce a parameter *θ ∈* [0, 1] that modifies both the initial resistant fractions and transition probabilities between sensitive and resistant phenotypes.

#### Pre-existing resistant fractions

The initial double-resistant fraction is defined as

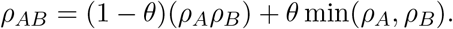

When *θ* = 0, double resistance follows the independence expectation. When *θ* = 1 and *ρ*_*A*_ = *ρ*_*B*_, all resistant cells are double resistant.

#### Transition probabilities

The probability of generating double resistance from a sensitive cell during division is defined as

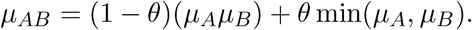

When *θ* = 0, double resistance arises only through independent acquisition of each single resistant phenotype. When *θ* = 1 and *µ*_*A*_ = *µ*_*B*_, transitions occur exclusively from *S* to *R*_*AB*_.

### 6.4 Simulation of experimental evolution

To simulate experimental resistance evolution under the two-drug model, we implemented two previously described modelling approaches (Whiting et al. 2025). We used (i) a hybrid phenotypic compartment model, which switches from stochastic jump dynamics to deterministic ODEs when compartment sizes exceed *N*_switch_, and (ii) a fully stochastic agent-based model in which each of the initial *N*_0_ cells is assigned a unique heritable barcode.

Simulations recapitulate the structure of long-term *in vitro* evolution experiments, including a shared drug-free expansion phase of duration *t*_exp_ following assignment of a barcode, sampling into *n*_rep_ replicate populations of size *N*_seed_, drug-specific treatment schedules, and optional passage bottlenecks at predetermined times. From each simulated experiment, for drug *X*, replicate *r* and passage *j*, we extracted the quantities used for inference: population sizes and passage times 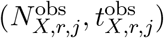 and barcode count vectors **c**_*X,r,j*_. Barcode counts were converted to relative frequencies and summarised using Hill diversity statistics of order *q* = 2, including within-replicate diversity 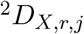,between-replicate diversity 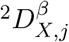, and between-drug diversity 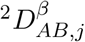, which quantify lineage relationships within and across replicates and drug conditions.

### 6.5 Bayesian inference framework

Model parameters were inferred using a two-stage simulation-based Bayesian framework that integrates population size trajectories with barcode diversity statistics. In the first stage, evolutionary parameters governing resistance to each drug were inferred independently. For a given drug *X ∈ {A, B}*, we inferred the parameter set Φ_*X*_ = *{ρ*_*X*_, *µ*_*X*_, *D*_*c,X*_, *κ*_*X*_, *Ψ*_*X*_*}*. Approximate Bayesian Computation with Sequential Monte Carlo (ABC–SMC) was first used to constrain parameter space based on observed population trajectories 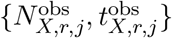 (Schäalte et al. 2022). To incorporate lineage information, we then applied Neural Posterior Estimation (NPE), a likelihood-free simulation-based inference method in which parameter samples from the ABC–SMC posterior were used to generate simulated diversity statistics via the agent-based model (Boelts et al. 2025). These statistics comprised within-replicate diversity 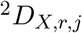 and between-replicate beta diversity 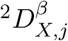 (order *q* = 2 Hill diversity). A neural density estimator was trained on pairs 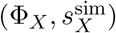 to approximate the posterior distribution 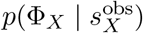, yielding independent posterior distributions for each drug.

In the second stage, cross-resistance strength was inferred using parameter draws from the independently inferred single-drug posteriors. Paired parameter sets (Φ_*A*_, Φ_*B*_) were sampled independently and combined with prior values of the strength of cross-resistance parameter *θ*. For each triplet (Φ_*A*_, Φ_*B*_, *θ*), the complete two-drug experiment (including the shared expansion phase of duration *t*_exp_ and replicate sampling structure) was simulated, and between-drug diversity statistics 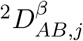 were computed per replicate. Neural Posterior Estimation was again used to approximate the posterior distribution of *θ*.

### 6.6 Estimating the strength of cross-resistance from synthetic data

To validate our inference framework, we simulated biologically plausible evolutionary scenarios with known values of the cross-resistance parameter *θ*. Using these synthetic datasets, we then applied our Bayesian inference procedure to assess its ability to recover the true underlying value of *θ*.

Cells were assigned a unique heritable barcode (*N*_0_ = 10^5^) before undergoing a shared expansion phase (*t*_exp_ = 8.0 days). For each drug condition (*Drug X ∈ {A, B}*), cells were seeded into replicate flasks (*n*_rep_ = 4) at a fixed seeding density (*N*_seed_ = 10^5^). Treatment was applied continuously, beginning one day after seeding (treat_on_ = [1.0]). Each replicate flask was simulated until either a maximum duration was reached (*t*_max_ = 100 days) or the population exceeded a maximum size (*N*_max_ = 10^7^). Flasks were grown for a single passage event (*J* = 1). For each drug, the final population size and simulation time 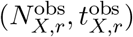 were recorded alongside the corresponding cell lineage (barcode) counts 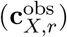. These data, simulated with known parameter values, produced ground-truth synthetic datasets used to benchmark our inference framework.

For the within-drug inference, for the first step each ABC–SMC run consisted of *G* = 10 generations with *n* = 2000 particles per generation and an acceptance threshold of *ϵ* = 0.01. For the second within-drug inference step, the NPE was trained using *n* = 500 simulations over *K* = 2 generations, yielding posterior distributions for each drug: 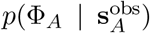 and 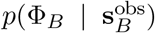. For the second between-drug stage, the NPE was trained using *n* = 500 simulations over *K* = 2 generations, yielding the posterior distribution for the strength of cross-resistance (*θ*).

### 6.7 Prior distributions

For single-drug inference, weakly informative priors were placed on all parameters.

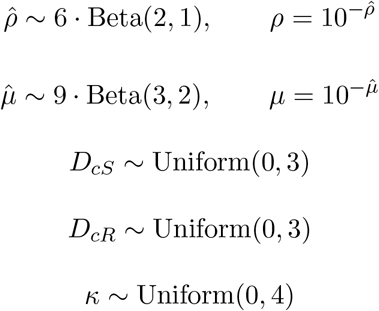

We fit separate drug-effect parameters for sensitive and resistant cells (*D*_*cS*_ and *D*_*cR*_). Under this formulation, differential drug sensitivity and resistance strength are captured via phenotype-specific drug kill strengths, with the (*D*_*c*_, *Ψ*) parameterisation recovered as the case *D*_*cS*_ = *D*_*c*_ and *D*_*cR*_ = (1 *− Ψ*)*D*_*c*_.

Parameters that control the abundance of resistance (*ρ* and *µ*) were defined on a log scale by sampling variables 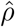 and 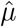 from scaled beta distributions and transforming via 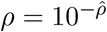 and 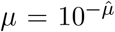. Given resistance is often assumed to be a rare event, this parameterisation concentrates prior mass on lower pre-existing frequencies and transition rates while allowing variation over several orders of magnitude.

For cross-resistance inference, the strength of cross-resistance *θ* was parameterised by sampling 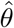 ~ Uniform(0, 3) and transforming via 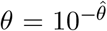. Because *θ* directly controls the initial abundance of double-resistant cells (which subsequently expand exponentially) this prior is also defined on a log scale and places most mass at low values, requiring strong evidence to support high levels of cross-resistance.

### 6.8 Estimating cross-resistance in ER^+^ breast cancer cell models treated with CDK4/6 inhibitors

We applied our cross-resistance inference framework to experimental data generated from three ER^+^ breast cancer cell models (MCF7, BT474, and T47D) treated with the CDK4/6 inhibitors palbociclib and abemaciclib (Mavrommati et al. 2025). For each cell model, cells were independently transduced with a high-complexity barcode library. As such, lineage identities are not comparable across models. Following barcoding, cells underwent a shared expansion step before being seeded into replicate flasks and treated in parallel with either palbociclib or abemaciclib under continuous exposure. Cell population sizes and barcode lineage abundances were recorded at the experimental endpoint.

Inference was performed independently for each cell model using fixed experimental parameters matched to the corresponding *in vitro* protocol. Initial barcoded population sizes were cell line–specific (*N*_0_ = 9.0 *×* 10^5^, 1.44 *×* 10^6^, and 3.9 *×* 10^6^ for MCF7, BT474, and T47D cells, respectively). Cells then underwent a two-stage expansion phase. The first expansion stage lasted *t*_exp_ = 10, 14, and 10 days for MCF7, BT474, and T47D cells, respectively, after which populations were bottlenecked and reseeded (*N*_seed_ = 3.0 *×* 10^7^, 4.0 *×* 10^7^, and 3.0 *×* 10^7^ cells). A second expansion stage of *t*_exp_ = 9 days followed for all cell lines, after which cells were seeded into *n*_rep_ = 3 replicate flasks per drug (*N*_seed_ = 1.5 *×* 10^6^, 3.0 *×* 10^6^, and 1.5 *×* 10^6^ cells for MCF7, BT474, and T47D cells, respectively). Treatment was initiated one day after seeding (treat on = [1.0]) and maintained throughout the experiment. Each replicate was simulated until either a drug-specific maximum duration was reached (*t*_max_ = 77 and 295 days for palbociclib and abemaciclib in MCF7 cells; 108 and 143 days in BT474 cells; 77 and 189 days in T47D cells) or the population exceeded a maximum size (*N*_max_ = 5*×*10^7^). Single-drug evolutionary parameters were first inferred separately for palbociclib and abemaciclib using population size trajectories and within-drug barcode diversity statistics. Cross-resistance inference was then performed by simulating the two drug experiments. This procedure yielded a posterior distribution for the cross-resistance parameter *θ* for each cell model, resulting in three independent estimates of CDK4/6 inhibitor cross-resistance corresponding to MCF7, BT474, and T47D cells.

### 6.9 Estimating cross-resistance in DLBCL treated with R-CHOP omponents

We applied our cross-resistance inference framework to a previously published dataset in which a diffuse large B-cell lymphoma (DLBCL) cell model (Pfeiffer cells) was evolved under exposure to components of the R-CHOP regimen(Adam C Palmer, Chidley, and Peter K Sorger 2019). In the original experiment, cells were subjected to random mutagenesis prior to treatment and exposed to short, intermittent drug schedules.

We analysed four drug conditions: 4-hydroperoxy-cyclophosphamide (4-H-Cyclo), Doxorubicin, Rituximab, and Vincristine. For each drug, we used the sequenced barcode count distributions and corresponding population size measurements reported at the experimental endpoint. Lineage identities were aligned within each treatment condition.

Experimental parameters specified by the original protocol, including the initial barcoded population size (*N*_0_ = 1.0 *×* 10^6^), the shared pre-treatment expansion period (*t*_exp_ = 12 days), the number of replicate populations per drug (*n*_rep_ = 3), and the maximum population size (*N*_max_ = 80 *×* 10^6^), were fixed prior to inference. Following expansion, cells were seeded at drug-specific densities (*N*_seed_ = 1.2 *×* 10^7^ for 4-hydroperoxy-cyclophosphamide, doxorubicin, and vincristine; 6 *×* 10^6^ for rituximab) and subjected to the treatment schedules defined by the experimental protocol. Treatment on/off windows were treat on = [0, 14] and treat off = [3, 17] days for 4-hydroperoxy-cyclophosphamide and doxorubicin, and treat on = [0, 7] and treat off = [3, 10] days for rituximab and vincristine. Replicate’s total population sizes were recorded at *t*_keep_ = 14 days for 4-hydroperoxy-cyclophosphamide and doxorubicin and *t*_keep_ = 7 days for rituximab and vincristine, with simulations terminated at drug-specific maximum durations (*t*_max_ = 28, 29, 14, and 17 days, respectively). All evolutionary parameters were inferred from the data using the procedure described above.

Because the experimental design involved four drugs, cross-resistance inference was performed pairwise between treatment conditions. For each drug pair, single-drug parameters were inferred independently. Conditional on these single-drug posteriors, cross-resistance inference was performed using between-drug barcode diversity statistics. This yielded separate posterior distributions for the cross-resistance parameter *θ* for each pairwise comparison.

### 6.10 Estimating cross-resistance in NSCLC treated with EGFR and MEK inhibitors

We next applied the framework to a previously published dataset in which a non-small cell lung cancer (NSCLC) cell model was evolved under exposure to an EGFR inhibitor (gefitinib) and a MEK inhibitor (trametinib) (Acar et al. 2020). For each drug condition, we used the reported sequenced barcode distributions and corresponding population size measurements from replicate treatment populations.

Experimental parameters specified by the original study including the initial population size (*N*_0_ = 1 *×* 10^6^), replicate number (*n*_rep_), treatment schedules (treat on = [7] and treat off = [1000] days for both drugs), the maximum population size (*N*_max_ = 5 *×* 10^7^), and drug-specific maximum durations (*t*_max_ = 35 days for gefitinib and 70 days for trametinib) were fixed prior to inference. Cells first underwent a two-stage expansion phase: an initial expansion lasting *t*_exp_ = 28 days, followed by a reseeding bottleneck (*N*_seed_ = 1.6 *×* 10^7^ cells) and a second expansion lasting *t*_exp_ = 14 days. Following this second expansion, cells were seeded into replicate drug flasks (*N*_seed_ = 1.5 *×* 10^7^ cells per replicate) prior to treatment. All evolutionary parameters were inferred using the two-stage inference framework described above. Single-drug parameters were first inferred independently for gefitinib and trametinib. Cross-resistance inference was performed by simulating paired drug experiments and computing between-drug diversity statistics, yielding a posterior distribution for *θ*.

### 6.11 Simulation-based evaluation of treatment switching and synthetic clinical trials

To estimate the clinical benefit of treatment switching, we simulated tumour dynamics using parameter sets either drawn from the joint posterior distribution inferred from experimental data or using pre-determined values. Each draw comprised drug-specific evolutionary parameters for Drug A and Drug B Φ_*A*_ and Φ_*B*_, together with the cross-resistance *θ ∈* [0, 1]. For a given draw, tumour growth under first-line therapy was simulated until the total population size *N* (*t*) first exceeded a predefined progression threshold *N*_*p*_. The time to reach this threshold under Drug A (or Drug B) is denoted *T*_*A*_ (or *T*_*B*_). At this timepoint the full phenotypic composition (*S, R*_*A*_, *R*_*B*_, *R*_*AB*_) was recorded and used as a common initial condition for two second-line branches: continued treatment (AA or BB) or treatment switching (AB or BA). All evolutionary parameters were held fixed and the same phenotypic composition was used for each branch, ensuring that divergence after progression arose solely from treatment history rather than stochastic variation prior to progression. Each branch was simulated until the total population size exceeded a second predefined threshold *N*_*cf*_, representing clinical failure, yielding

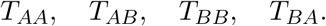

If extinction occurred before reaching *N*_*cf*_, the failure time was set equal to the maximum permitted simulation time (*t*_max_). The clinical benefit of switching from A to B was defined as

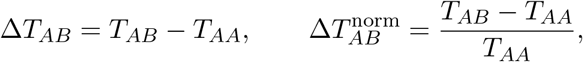

with an analogous definition for switching from B to A. To isolate the specific contribution of cross resistance, simulations could be repeated with *θ* fixed to its theoretical minimum (0.0) and maximum (1.0) values whilst allowing all other parameters to vary according to the joint posterior distribution. This defines a cross-resistance window within which the predicted benefit of switching can be attributed to cross resistance itself rather than to other drug-specific evolutionary behaviours.

To simulate synthetic clinical trials, each posterior distribution for a given cell model and drug comparison was interpreted as defining a “patient archetype”, and synthetic cohorts were generated by repeated sampling from the corresponding joint posterior. For each synthetic patient we simulated the relevant treatment arms (e.g. AA versus AB, or BB versus BA) and computed time-to-progression (TTP) using criteria applied to the simulated total population size trajectory. Progression was assessed only after the first crossing of a reference threshold *N*_prog_. Thereafter, observations were evaluated on a discrete recording schedule with interval Δ*t* = 14 days with a random phase offset uniformly distributed in [0, 14] days. Progression was called at the first recorded timepoint satisfying both

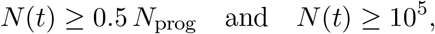

ensuring that the tumour reached a minimum ‘detection threshold’. TTP was reported in months. Kaplan–Meier curves were constructed for each drug switching arm.

---

**Extended Data Fig. 1.**
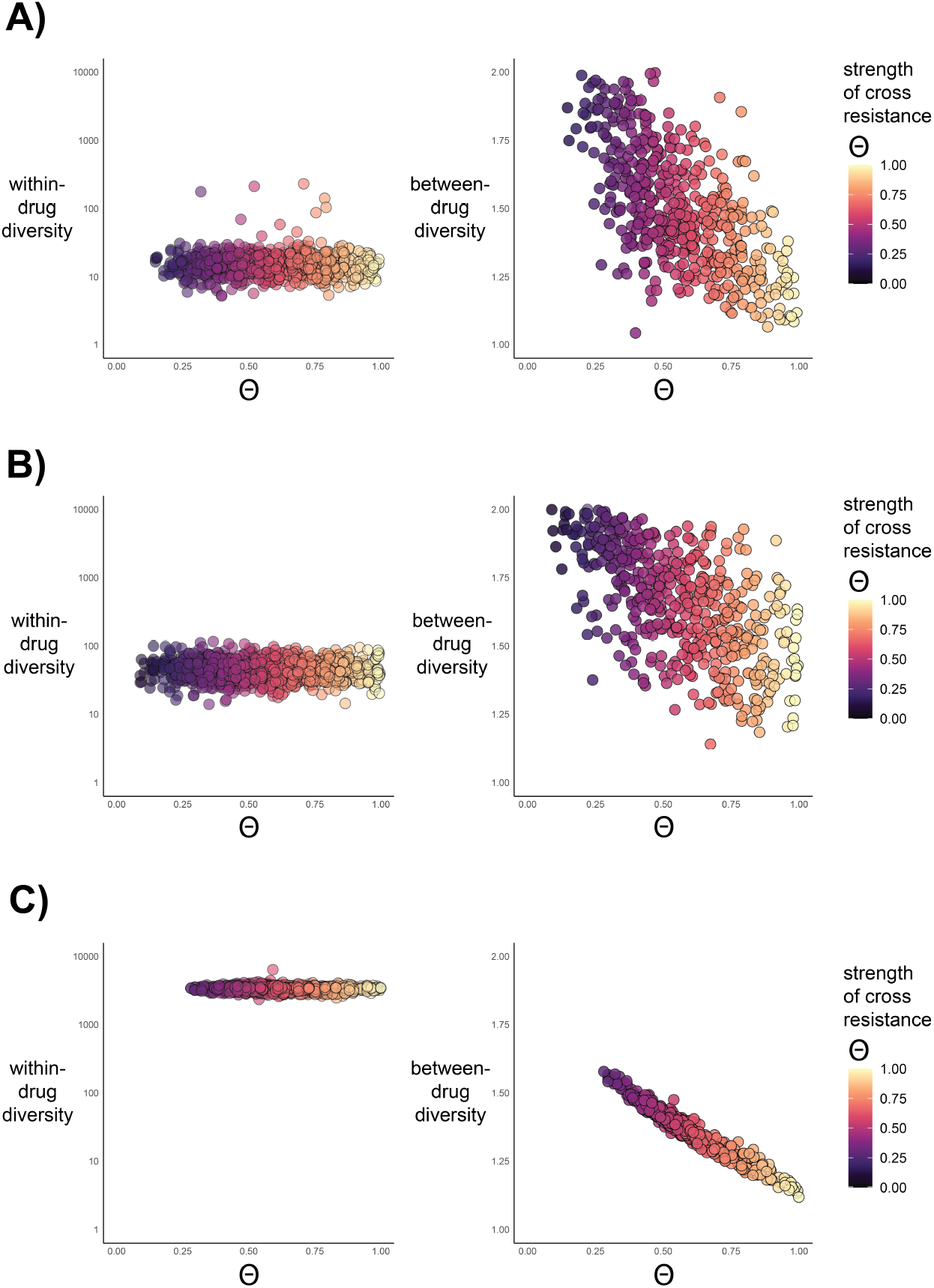
Within- and between-drug diversity statistics across simulated levels of cross resistance. Within-drug (left panels) and between-drug (right panels) diversity statistics plotted as a function of the cross resistance parameter (*θ*), calculated from simulated datasets (n = 500 per simulation). Panels show scenarios with (A) low, (B) intermediate, and (C) high levels of resistance to both drugs: *Drug A* and *Drug B*.

**Extended Data Fig. 2.**
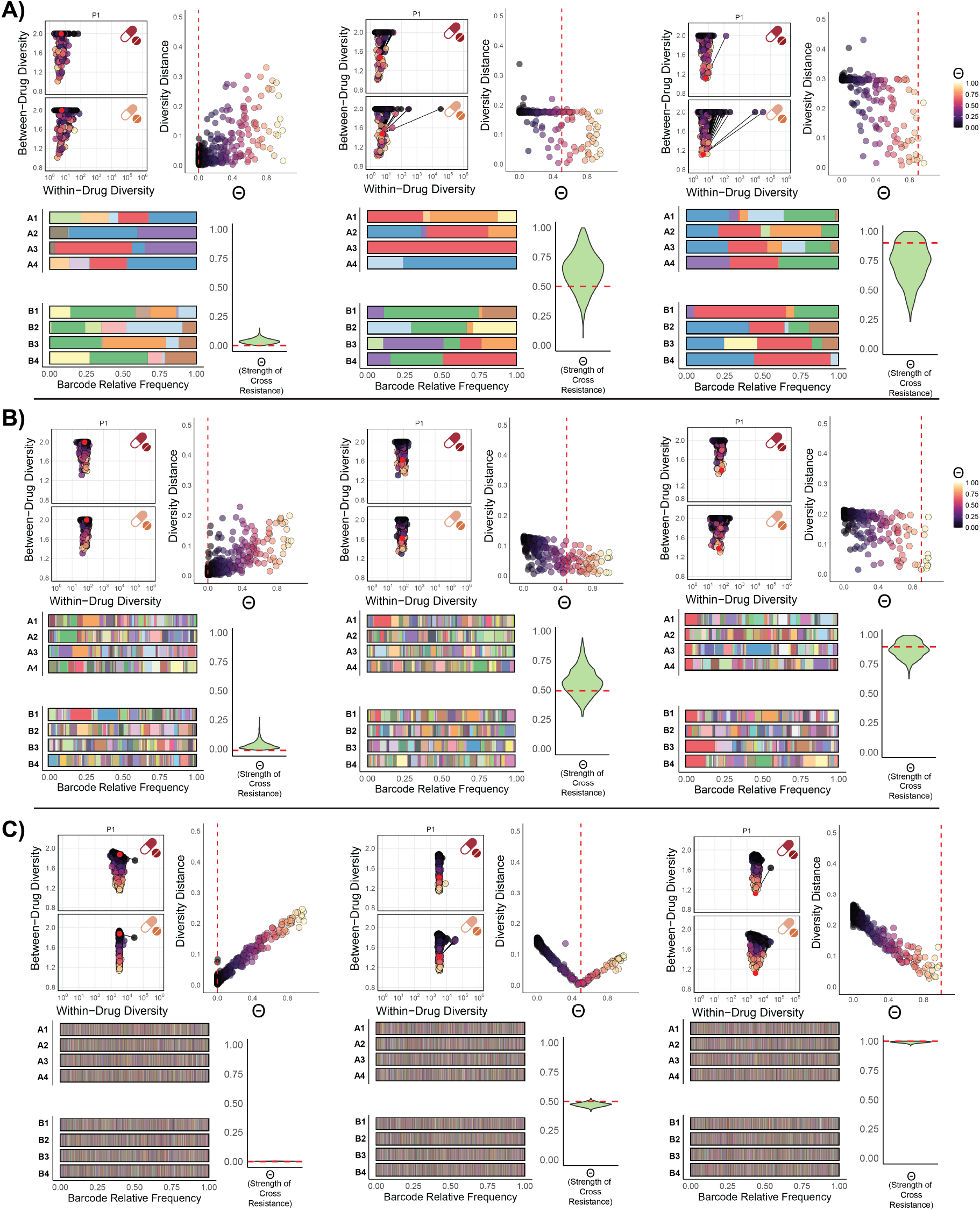
Inferred cross resistance across a range of evolutionary scenarios. Within- and between-drug diversity statistics used during the cross-resistance inference for *Drug A* and *Drug B*. Within each of panels (A–C), the within-drug parameters are held constant while the strength of cross resistance varies. Ground-truth statistics and the true cross resistance strength (*θ*) are shown as red points and dashed lines. Simulated barcode distributions and inferred cross-resistance values are shown for three evolutionary scenarios (A-C) for low (left), intermediate (middle) and high (right) levels of *θ* and low (A), intermediate (B) and high (C) levels of resistance. A subset of within- and between-drug diversity values (n=200 simulations per parameter set) are shown following the first generation of the cross resistance inference workflow. Barcode distributions are the simulated ground-truth sets used for inference.

**Extended Data Fig. 3.**
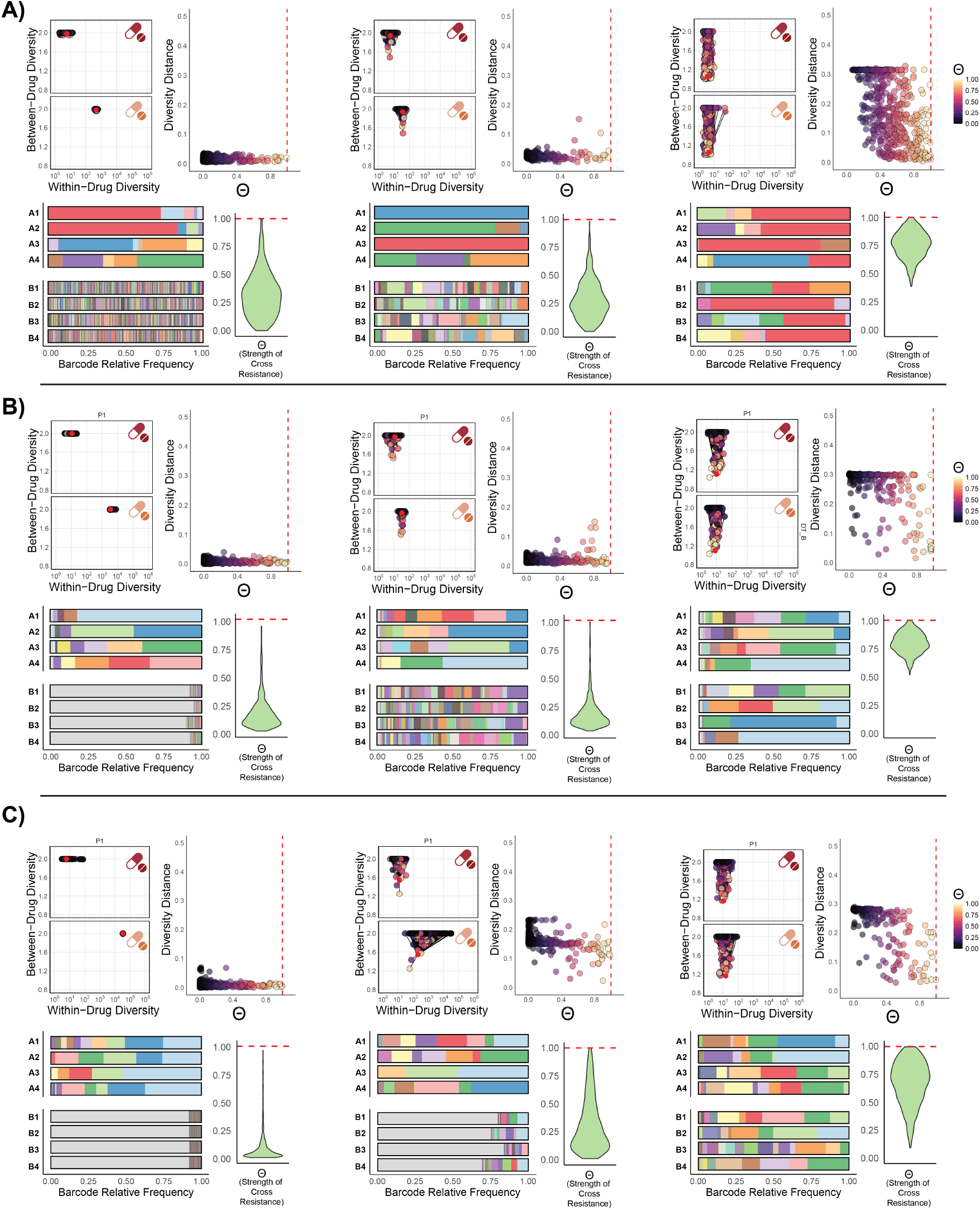
Cross-resistance inference under asymmetric single-drug evolutionary dynamics. Within- and between-drug diversity statistics used during the cross-resistance inference for *Drug A* and *Drug B*. Within each of panels (A-C), the strength of cross resistnace was held constant while different individual within-drug behaviours were allowed to differ between drugs. Ground-truth statistics and the true cross-resistance strength (*θ*) are indicated by red points and dashed lines. Simulated barcode distributions and inferred cross-resistance values are shown for three classes of asymmetry: (A) differences in the pre-existing resistant fraction (*ρ*), (B) differences in the transition probability from sensitive to resistant (*µ*), and (C) differences in drug effect strength (*D*_*c*_). Within each panel, moving from left to right corresponds to increasing levels of asymmetry between the two drugs. A subset of within- and between-drug diversity values (n=200 simulations per parameter set) are shown following the first generation of the cross resistance inference workflow. Barcode distributions are the simulated ground-truth sets used for inference.

**Extended Data Fig. 4.**
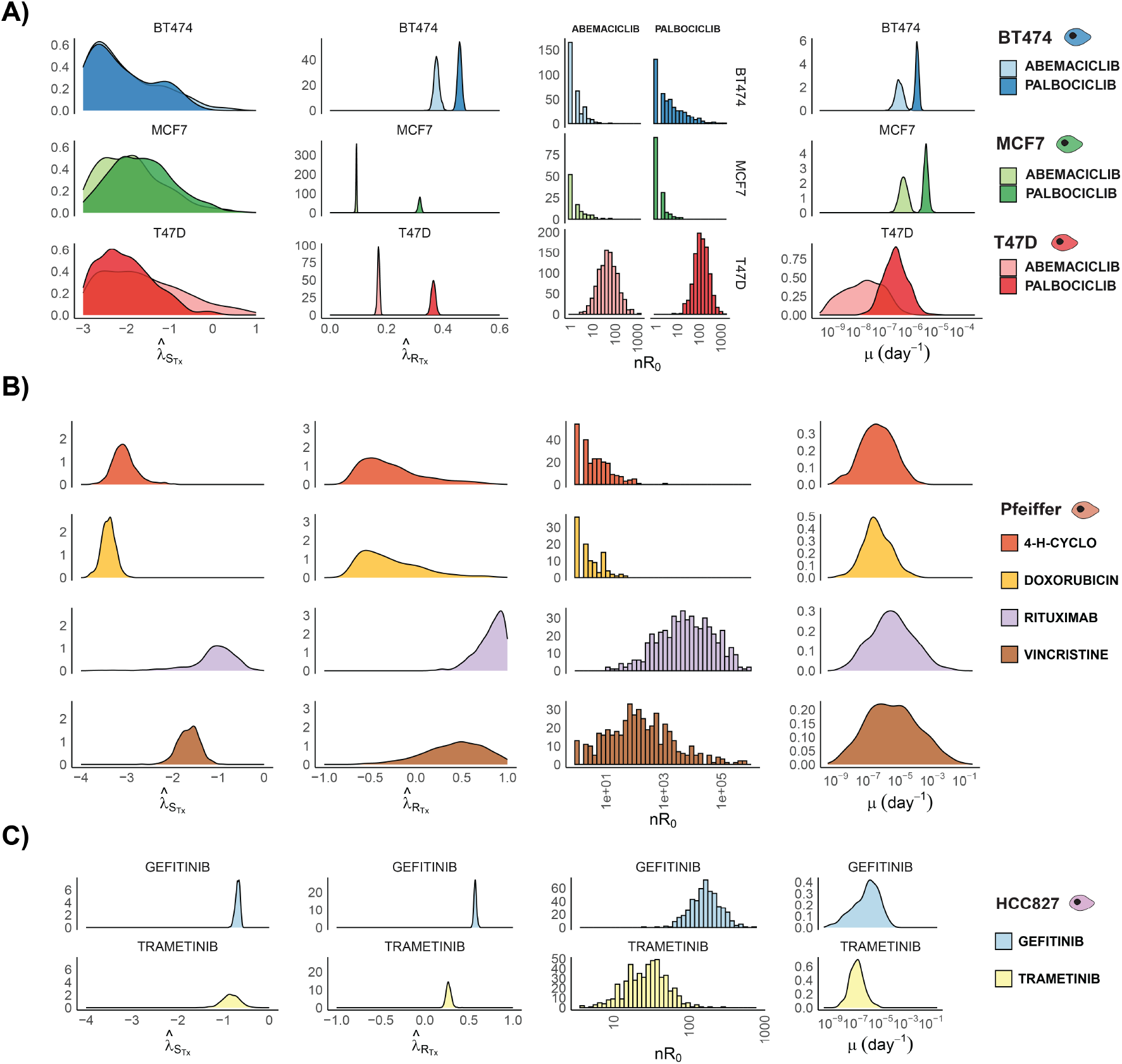
Single-drug behaviour posterior estimates from experimental data. Posterior distributions of the four evolutionary behaviours inferred using the Bayesian inference framework: normalised sensitive and resistant phenotype growth rates during treatment 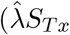 and 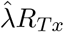, respectively), the pre-existing number of resistant cells at the time of barcoding (*nR*_0_) and the phenotypic transition rate from sensitive to resistant (*µ*). Estimates are derived using data generated from (A), lymphoma (B) and lung cancer (C) resistance evolution experiments.

**Extended Data Fig. 5.**
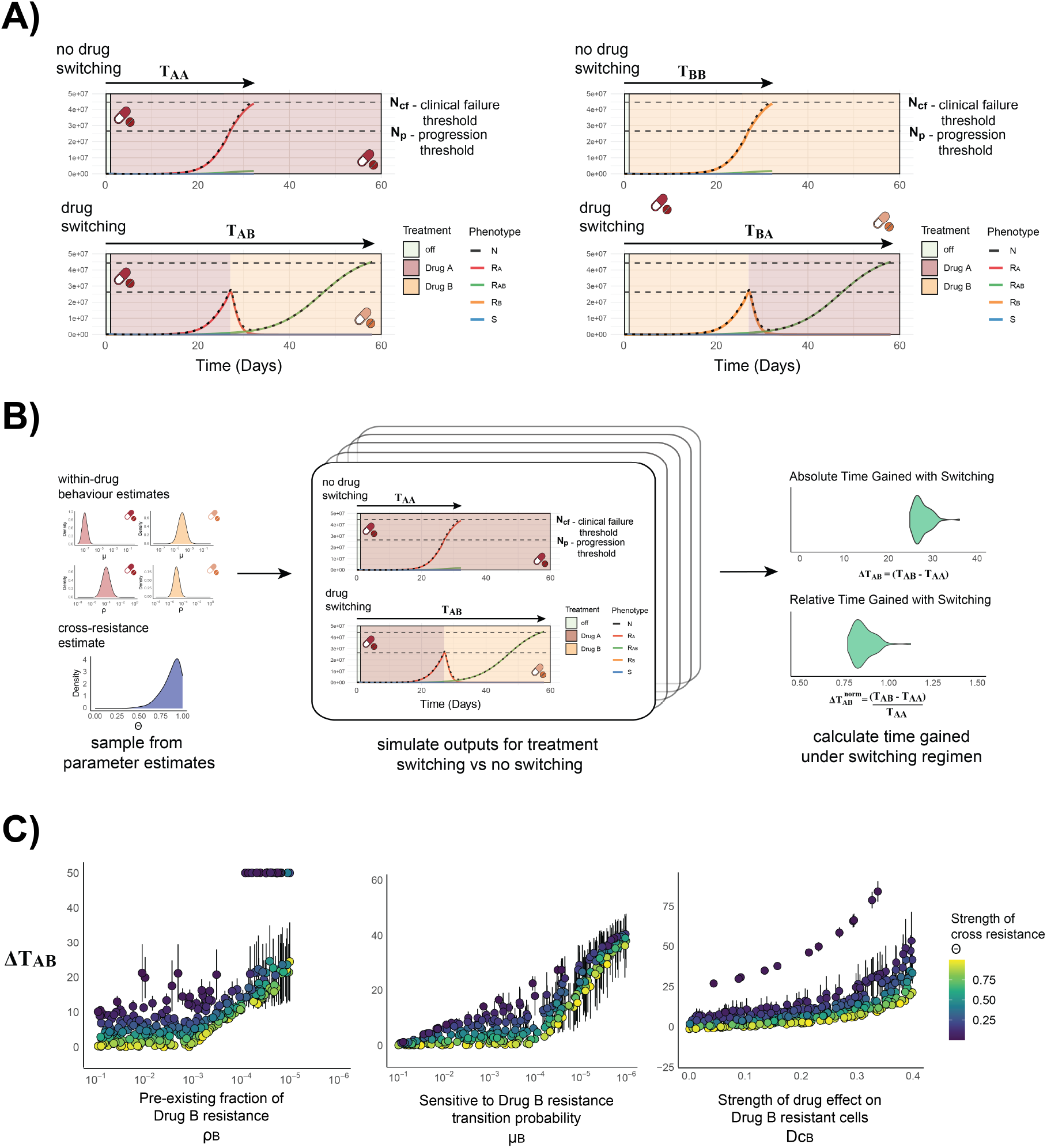
A simulation framework for comparing treatment sequence regimens given within- and between-drug behaviours. (A) Schematic illustrating the measured time during simulations until progression (*N*_*p*_) and clinical failure (*N*_*cf*_) under different treatment sequences compared with no switching. (B) Workflow for using inferred within-drug behaviours and cross-resistance estimates to calculate the absolute and relative time gained under a treatment-switching regimen. (C) Simulation results (n = 400 simulations, 50 replicates per simulation) illustrating the relationship between parameters governing single-drug resistance to *Drug B* (*ρ*_*B*_, *µ*_*B*_, *D*_*c,B*_) and the predicted benefit of switching from *Drug A* to *Drug B* compared with continuation of *Drug A*, measured in days (Δ*T*_*AB*_). Points indicate posterior mean parameter values, shading denotes the strength of cross resistance (*θ*) and error bars indicate *±* one standard deviation.

**Extended Data Fig. 6.**
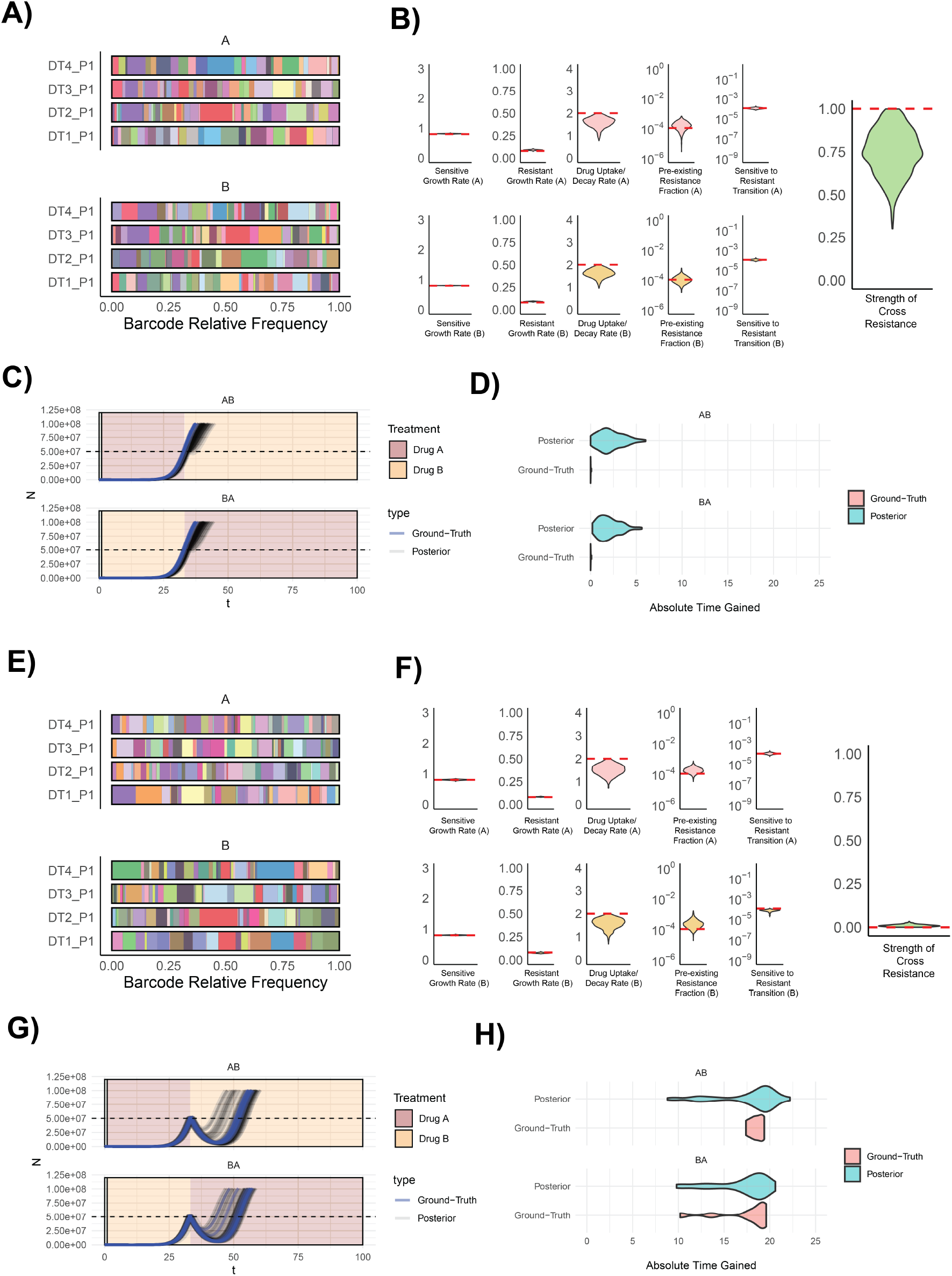
The inference framework can accurately predict the population dynamics under treatment-switching strategies. Simulated barcode distributions (A,E) and inferred within-drug behaviours and between-drug cross-resistance parameters (B,F) obtained using the Bayesian inference framework. Ground-truth values are highlighted (red dashed lines). Ground-truth parameter sets are shown alongside population size dynamics simulated using posterior parameter estimates under two treatment-switching regimens (n=50 simulations per condition) (C,G). Comparisons of the ground-truth and posterior estimates of the absolute time gained under each treatment strategy are shown in (D,H). Time gained is calculated as the delay in reaching the clinical failure threshold (*N*_*cf*_) relative to the corresponding no-switching condition (not shown).

